# Changes of chromosomal architecture before establishment of chromosome territories revealed by recurrence plot reconstruction

**DOI:** 10.1101/2021.05.20.444916

**Authors:** Yuki Kitanishi, Hiroki Sugishita, Yukiko Gotoh, Yoshito Hirata

**Author notes:** Corresponding authors: Hiroki Sugishita, International Research Center for Neurointelligence (WPI-IRCN), The University of Tokyo, 7-3-1 Hongo, Bunkyo-ku, Tokyo 113-0032, Japan, Yoshito Hirata, Faculty of Engineering, Information and Systems, University of Tsukuba, 1-1-1 Tennodai, Tsukuba, Ibaraki 305-8573, Japan.

## Abstract

The chromatin conformation capture-related methods such as Hi-C have improved our understanding of nuclear architecture and organization in recent years. However, reconstruction of nuclear architecture from single-cell Hi-C (scHi-C) data is challenging due to limited information of DNA contacts obtained from a single cell. We have previously developed the Recurrence Plot-Based Reconstruction (RPR) method for reconstructing three dimensional (3D) genomic structure from Hi-C data of single haploid cells (1) and diploid cells (2). This algorithm is based on a recurrence plot, a tool of nonlinear time-series analysis for visualizing patterns within a time series (3, 4), and enables the reconstruction of a unique 3D chromosome architecture even from low-coverage DNA contact information. Here we used the RPR method to analyzing published scHi-C data of diploid cells derived from early-stage F1 hybrid embryos (5) as a proof-of-concept for understanding of global developmental changes in chromosomal architecture of early stage embryos. We found that paternal and maternal chromosomes become gradually intermingled from 1 cell to 64 cell stage, and that discrete chromosome territories are largely established between 8 cell and 64 cell stages. We also observed Rabl-like polarization of chromosomes from the 2-to 8-cell stages, but this polarization becomes mostly dissolved by the 64-cell stage. Rabl-like chromosome polarization precedes rod-like extension and parallel alignment of chromosomes, implicating the role of Rabl-like polarization in effective mixing of chromosomes before establishing chromosome territories. We also found cell-to-cell variability in chromatin configuration. A combination of scHi-C and RPR analyses can depict features of the 3D chromatin architecture of individual cells at different developmental stages during early embryogenesis.

## Introduction

Spatial organization of the mammalian genome plays a key role in transcription, RNA processing, and DNA replication (6–9). The chromatin is hierarchically organized into local loops, into different scales of domains and compartments, and into chromosome territories (CTs) (10–12). Hierarchical organization of chromatin is a foundation for gene regulation and cell function. Classically, spatial localization of genomic region-of-interest in the nucleus has been elucidated with DNA-fluorescence in situ hybridization (FISH) (13).However, although recent endeavors have increased the number of genomic loci to be analyzed (14, 15), FISH-based methods can analyze only targeted genomic locus-of-interest. By contrast, Hi-C, a derivative of chromosome conformation capture(16), can elucidate nontargeted and genome-wide pair-wise DNA contacts by sequencing after proximity-dependent ligation (17). Application of Hi-C to bulk cell populations (bulk Hi-C) has revealed principles of chromatin organization, such as organization of chromatin into topology-associated domains and A/B compartments(17–20). Because bulk Hi-C per se cannot map regions in relation to the global nuclear geography (the 3D genome atlas), bulk Hi-C is used with other imaging and epigenetic analyses (such as DNA FISH and Lamin-DamID) to map regions in the 3D genome atlas and to understand how chromatin architectures are coordinated in the nucleus (21–28). Bulk Hi-C can provide information about contacts between genomic regions on the basis of proximity ligation, but this method is limited to measuring simultaneous contacts between two genomic regions and thus cannot provide information about how multiple genomic regions are simultaneously organized in the nucleus(29). Other methods can simultaneously identify contacts involving multiple genomic regions. These methods include genome architecture mapping (GAM, which is based on ultrathin cryo-sectioning, DNA sequencing, and the mathematical model SLICE) and split-pool recognition of interactions by tag extension (SPRITE, which identifies nearby genomic regions by barcoding during split-pool cycles) (30) (31). However, these methods cannot generate a 3D genomic atlas at the single-cell level because they estimate average contacts from an ensemble of cells.

The 3D architecture of chromatin changes dramatically during cell differentiation and during cell state transitions. Even between cells of the same type and state, chromatin architecture is highly dynamic and heterogenous. It is thus crucial to investigate the 3D genome atlas at a single cell level. Single-cell Hi-C (scHi-C) and in situ genome sequencing (IGS) have been developed to construct 3D genome atlases for single cells in a nontargeted and genome-wide manner. IGS is a method to spatially map genome-wide paired-end sequence reads to their endogenous locations within intact cells. scHi-C (a modified version of widely used Hi-C) has been applied to both haploid and diploid cells (27, 32–36). In diploid scHi-C, allele-specific single nucleotide variation (SNV) are used to differentiate paternal and maternal alleles. However, reconstruction of 3D nuclear architecture from scHi-C data has been challenging due to a limited number of DNA contacts recovered from a single cell as well as the limited frequency of SNVs within the genome. Tan et al. has developed a modified scHi-C method for diploid cells called Dip-C that combines a transposon-based whole genome amplification procedure and an algorithm for imputation of the two chromosome haplotypes linked by each contact (35), although this algorithm contains some imputations that are not necessarily proven mathematically. We reported another algorithm, the Recurrence Plot-Based Reconstruction (RPR) method, that reconstructs the 3D chromosome structure from scHi-C data of either haploid (1) or diploid cells (2). This method is based on a recurrence plot (a tool of nonlinear time-series analysis for visualizing patterns within a time series) (1) and enables a unique 3D chromosome architecture to be reconstructed from even low-coverage DNA-contact information. The RPR method has been mathematically proven (1, 37, 38). In contrast to Dip-C’s algorithm which assumes that paternal and maternal alleles are not spatially close to each other and do not have similar shapes, RPR algorithm only assumes that consecutive genomic regions are neighbors (1, 2).

As a proof-of-concept, we analyzed 3D nuclear architectures of cells at early preimplantation stages after fertilization with the use of RPR method and a set of scHi-C data publicly available(5), given that chromatins at these stages undergo dramatic changes as development proceeds (39). Our results confirm some of the observations obtained with FISH and IGS, such as gradual mixing of paternal and maternal alleles during progression from the zygote to the 64-cell stage. Our analyses also revealed that discrete (localized) CTs are established between 8-cell and 64-cell stages, and suggested potential roles of Rabl-like polarization and rod-like extension of chromosomal structure in CT establishment.

## Results

### Application of the RPR algorithm to scHi-C data of early-stage mouse embryos

We applied the RPR method to analyze scHi-C data from diploid cells of early-stage mouse embryos (1-cell stage [n = 75]; 2-cell stage embryos [n = 109]; 4-cell stage [n = 121]; 8-cell stage [n = 125]; and 64-cell stage [n = 201]) published by Collombet et al. (2020) (5). The data were obtained from highly polymorphic F_1_-hybrid embryos, generated by crossing female *Mus musculus domesticus* (C57Bl/6J) with male *Mus musculus castaneus* (CAST/EiJ) (Collombet et al., NCBI GSE129029). The paternal and maternal alleles were distinguished by SNVs between these genetic backgrounds: 22.6% (1-cell), 24.4% (2-cell) 21.2% (4-cell), 23.7% (8-cell) and 22.9% (64-cell) of genomic reads were uniquely assigned to the maternal (C57Bl/6J) genome, whereas 14.6% (1-cell), 18.0% (2-cell) 16.7% (4-cell), 17.9% (8-cell) and 17.7% (64-cell) of genomic reads were uniquely assigned to the paternal (CAST/EiJ) genome (Ext. Fig. 1b). For performing RPR, we divided the genome into 1 Mb segments and used contact information of genomic segment pairs in which both segments harbored at least one SNV. A summary of the RPR method is as follows (Fig. 1a) (2). First, we constructed a weighted network from a contact map obtained from a single diploid cell Hi-C dataset. In this network, each node corresponds to a segment of an allele-distinguished chromosome. We connected two nodes if there is a contact between these corresponding chromosomal segments both of which have their alleles identified with SNVs.

**Figure 1:**
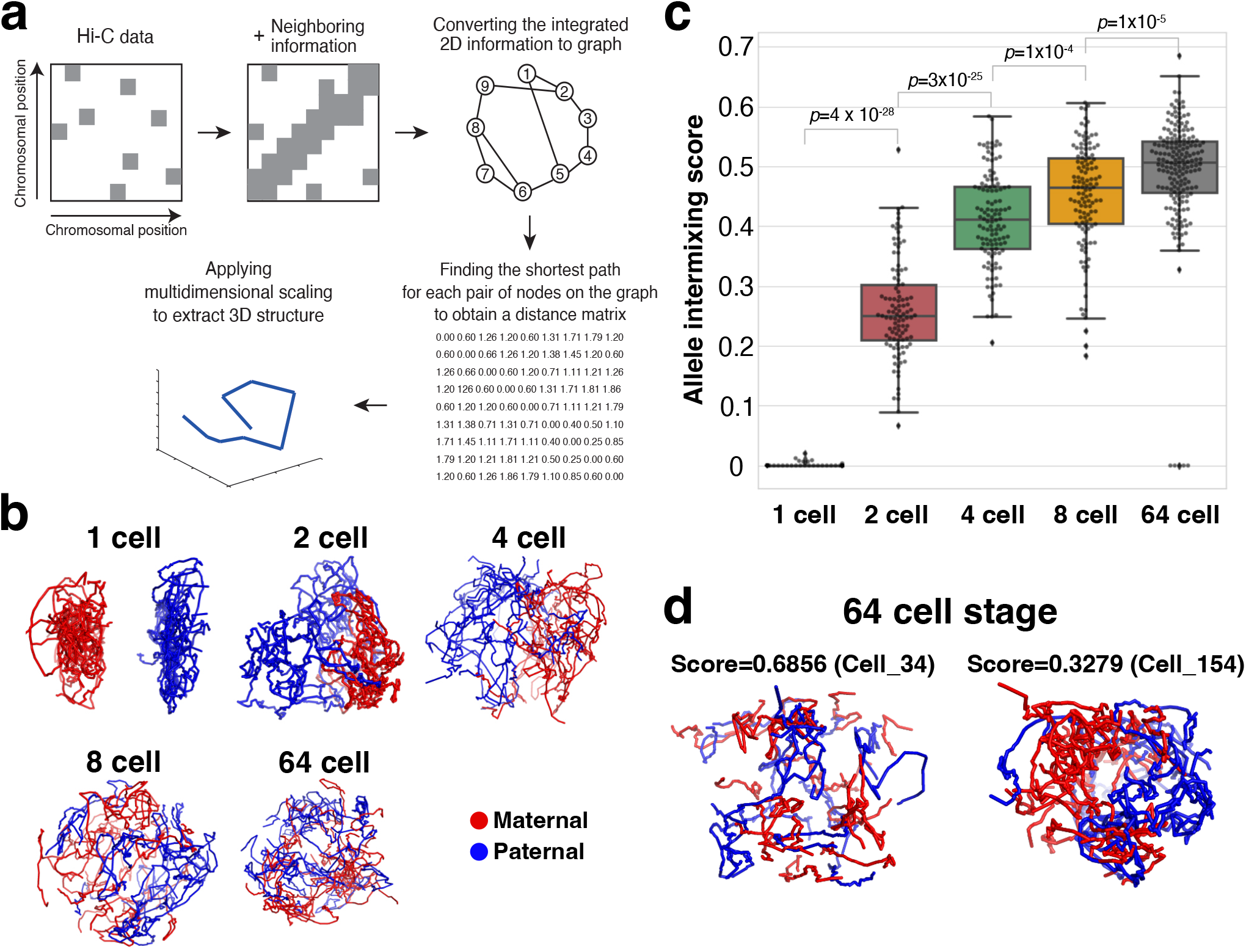
Spatial intermixing of paternal and maternal genomes during early stage development. **a**. Schematic illustration of the recurrence plot-based reconstruction (RPR). **b**. Three-dimensional reconstruction of representative single nuclei at different developmental stages. Paternal and maternal alleles are labeled in different colors (maternal, red; paternal, blue). **c**. Box plots showing the paternal-maternal allele intermixing score at different developmental stages. Each dot indicates the score of individual cells. The p value was calculated by Mann Whitney U test with Bonferroni correction. **d**. Three-dimensional reconstruction of representative single nuclei at 64 cell stage. Paternal and maternal alleles are labeled in different colors (maternal, red; paternal, blue), as in **b**. Note that the cell in the right side shows less intermixing of parental alleles than that in the left side.

We also connected two nodes if they are on the same allele of the same chromosome within 10 Mb. Let i and j denote a node in the network, respectively. Then, we assigned the weight for the edge between nodes i and j by using the portion of common nodes against the nodes which are connected with either i, j or both (this definition coincides with that of the Jaccard coefficient (40). More exactly, the weight is the portion of common nodes subtracted from 1. Then, this weight can be regarded as the local distance between two corresponding segments i and j. The latter parts are similar to Lesne et al., Hirata et al., and Tenenbaum et al. (41–43): we calculated the shortest distances between every pair of nodes in the network, followed by applying the classical multidimensional scaling to find a set of point arrangements preserving the shortest distances so that top three components become our 3D reconstruction of chromosomes. The strength of our method is that (i) we can reconstruct local structures more finely by reconstructing the local distances by the Jaccard coefficient appropriately compared with that of Lesne et al. (43), (ii) we can apply the method with sparser data because the proportion of common nodes is a robust statistic under such conditions, and (iii) we can use the information of pairs of chromosome segments in the Hi-C data, in which at least one of the segments does not harbor a SNV and thus were not used for 3D reconstruction, to evaluate the accuracy of reconstruction.

In this study, we defined the inaccuracy score for each reconstructed point by using the root mean square distance between each reconstructed point and the spatial average of its corresponding neighbors (the contact pairs found in the Hi-C data) (Ext. Fig. 1a). Here, the neighbors were defined as 1 Mb segments containing a contact pair or some. The more accurate the reconstruction is, the closer the neighbors are expected to be to the reconstructed point in the 3D genome atlas. With the RPR method, the center of the nucleus tends to be more accurate compared to the peripheral (Ext. Fig. 1c). The overall results indicate that a majority of reads, even those with no overlapping SNV, to be assigned to the predicted spatial location through co-localization with haplotype-resolved reads in the same location. However, some cells had relatively high average inaccuracy scores (with greater standard deviations [*S*.*D*.]; Ext. Fig. 1d,1e); these cells also tended to show irregularly shaped nuclei (for example, see Fig. 1d, left). We thus excluded the cells with high inaccuracy score (*S*.*D*. > 0.15) from the following analyses (Ext. Fig. 1f). We found no significant correlation between the inaccuracy score and the number of contact pairs detected by Hi-C (Ext. Fig. 1g), suggesting that the high inaccuracy score is not solely due to the few reads of Hi-C contact pairs.

### Spatial intermixing of paternal and maternal genomes during early-stage development

At the 1-cell stage, the parental genomes are separate, each in a haploid pronucleus. The parental genomes mix after this stage, but the time and extent of this mixing has been somewhat controversial (44, 45). A DNA-FISH analysis that separately visualized parental centromeres in mouse interspecific hybrids (as well as an analysis of sperm-specific BrdU labeling) has shown that parental genomes are topologically separate until the 4-cell stage and then gradually mix(44). By contrast, a recent IGS analysis has shown that parental genomes mix from 1-to 4-cell stages(45). We thus investigated the extent of mixing of the parental genomes based on the RPR of 3D genomic architecture. We observed the parental genomes to gradually mix from the 1-to 64-cell stages (Fig. 1b). To quantitively measure genome mixing, we first calculated the distance of all pairs of 1 Mb genomic segments. We then defined proximity pairs (two closely located genomic segments) as the top (nearest) 5% of all pairs (Extended Figure 2). To focus on inter-chromosomal arrangements, we only used the information about those genomic segments that had at least one proximity pair with a genomic segment outside of its own chromosome. We then calculated the fraction of proximity pairs with genomic segments containing different parental alleles (“the intermixing score”) in each cell. This fraction was close to zero at the 1-cell stage (as expected) but gradually increased as the zygote developed to the 64-cell stage (Fig. 1b). The mean intermixing score significantly increased between 1- and 2-cell stages; 2- and 4-cell stages; 4- and 8-cell stages; and 8- and 64-cell stages (Mann-Whitney U-test with Bonferroni correction, *p* < 10^−27^, *p* < 10^−24^, *p* < 10^−3^, *p* < 10^−4^) (Fig. 1c). This indicated that parental genome mixing starts at the 2-cell stage, consistent with the result from IGS analysis (45). Moreover, our results quantitatively demonstrate that parental genomes still significantly mix after 4 cell stage, as shown by the DNA-FISH analysis. Given that the intermixing score became close to 0.5 at the 64-cell stage (Fig. 1c), chromosomes appear to be positioned irrespectively to their parental alleles by this stage. Interestingly, however, the extent of chromosome mixing varies among cells at all stages (see examples of 64-cell stage mixing, Fig. 1d), suggesting cell-to-cell variability of chromosome organization, which has previously been proposed (44).

**Figure 2:**
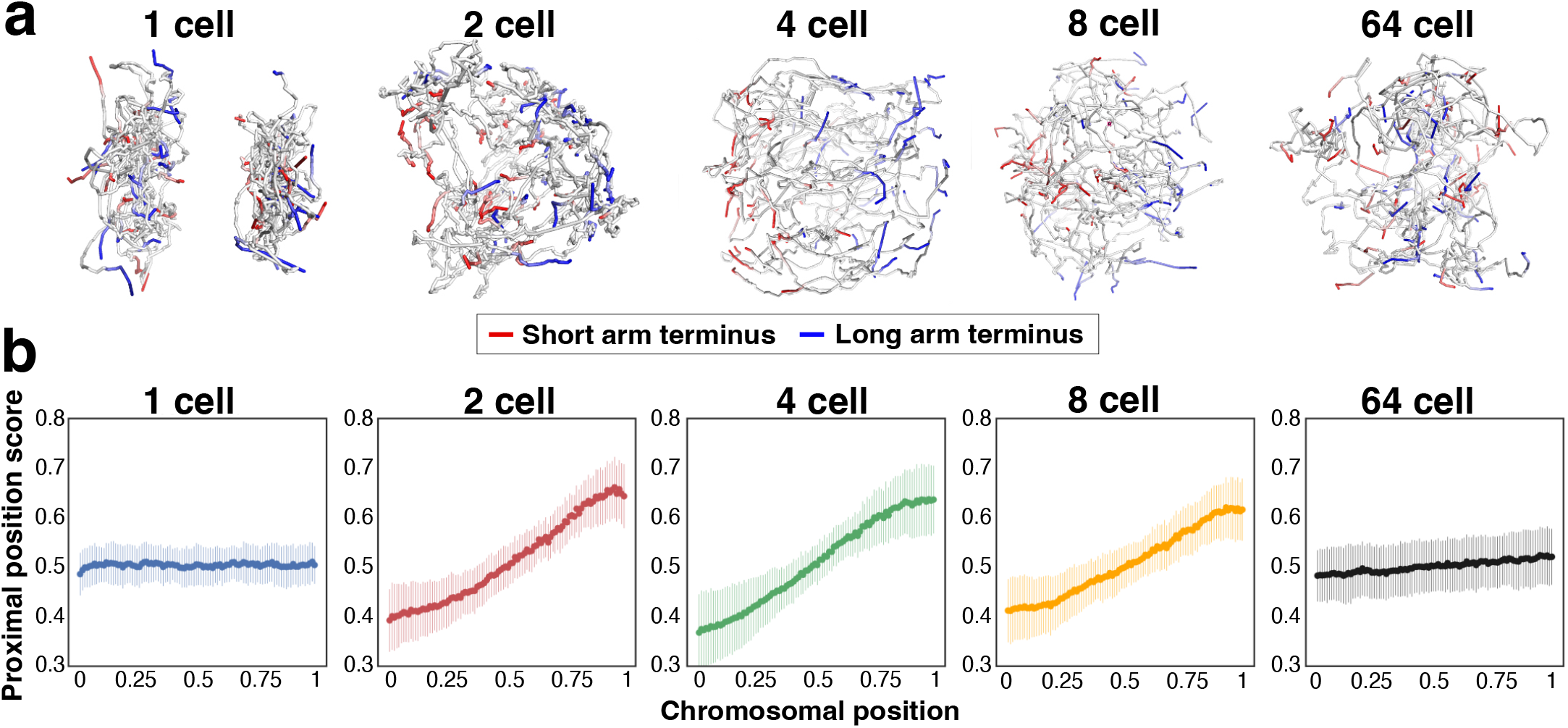
Formation and dissolvement of Rabl-like configuration during early-stage development. **a**. Three-dimensional reconstruction of representative single nuclei at different developmental stages. The terminus of the short arm and that of the long arm are highlighted in red and blue, respectively. **b**. The mean chromosomal position (the short arm terminus = 0, the long arm terminus = 1) of proximity pairs of each genomic segment (“Proximal Position Score”; y-axis). The chromosomal position of segment along the chromosomal axis (x-axis, the short arm terminus = 0, the long arm terminus = 1, all chromosomes were divided). s.d. for each segment of the averaged chromosome is shown.

### Formation and dissolvement of Rabl-like configuration during early-stage development

Mouse chromosomes are acrocentric. In an early-stage mouse embryo, chromosomes are organized into a Rabl-like, polarized configuration: centromeres and their nearby telomeres are confined to one part of the nucleus while the remaining telomeres cluster in the other part (45–48). The extent of this polarization at each developmental stage has been controversial. A 3D FISH analysis of centromeric, pericentromeric, and telomeric DNA sequences showed that polarity levels were rather constant from 2-to 32-cell stages (48). However, an IGS analysis showed that polarity increased from 1-to 4-cell stages (45). We thus measured chromosomal polarity along the centromere–distal telomere axis at each stage in our analysis. After labeling the short-arm and long-arm termini of each chromosome in the reconstructed 3D map, the polarization of chromosomes was visible from the 2-to 8-cell stages (Fig. 2a). To quantitatively analyze the extent of polarization, we calculated the mean position along the chromosomal axis (the short-arm terminus = 0, the long-arm terminus = 1) of proximity pairs for each genomic segment (but excluded proximity pairs comprising genomic segments within the same chromosome, as shown in Fig. 1c). We found a greater correlation (between the position of each genomic segment along the chromosomal axis and the mean chromosomal position of its proximity pairs) in 2-, 4-, and 8-cell stage embryos than in zygotes and 64-cell stage embryos (Fig. 2b). This indicates that chromosomes become polarized in a Rabl-like configuration from zygote-to 2-cell stages (as reported by Payne et al)(45) but this configuration becomes mostly dissolved from 8-to 64-cell stages. This dissolvement appears to start between the 4- and 8-cell stages, because the angle of the correlation curve becomes shallower at these stages (Fig. 2b). Our RPR analysis of scHi-C data thus indicated that polarity increases from 1-to 4-cell stages (this part is consistent with IGS results (45)), that polarity may start to decrease from 4-to 8-cell stage, but rather dramatically decreases between 8- and 64-cell stages.

### Establishment of discrete chromosome territories during early-stage development

Within the nucleus, each chromosome is confined to a discrete, patch-like region called a chromosome territory (CT) (10–12). The spatial organization of CTs is not random and critical for gene regulation and genome stability (12). It remains unclear, however, when discrete (patch-like) CTs emerge during early-stage development. We thus evaluated the extent of intra-chromosomal interactions (compared to that of inter-chromosomal interactions) at each stage by using the 3D genomic architecture based on the RPR method. We again defined proximity pairs as the top (nearest) 5% of all pairs and calculated the fraction of proximity pairs in which the genomic segments were on the same chromosome in each cell (“the CT score”). The CT score slightly increased from 1-to 8-cell stages and greatly increased from 8-to 64-cell stages (Fig. 3a and 3b). This suggests that discrete CTs are largely established between the 8- and 64-cell stages. The increase of the CT score between the 8- and 64-cell stages was found in all chromosomes but to a lesser extent for shorter chromosomes especially for chromosome 19 (Fig. 3c), probably because of their already packed (nonelongated) configuration at the 8-cell stage.

**Figure 3:**
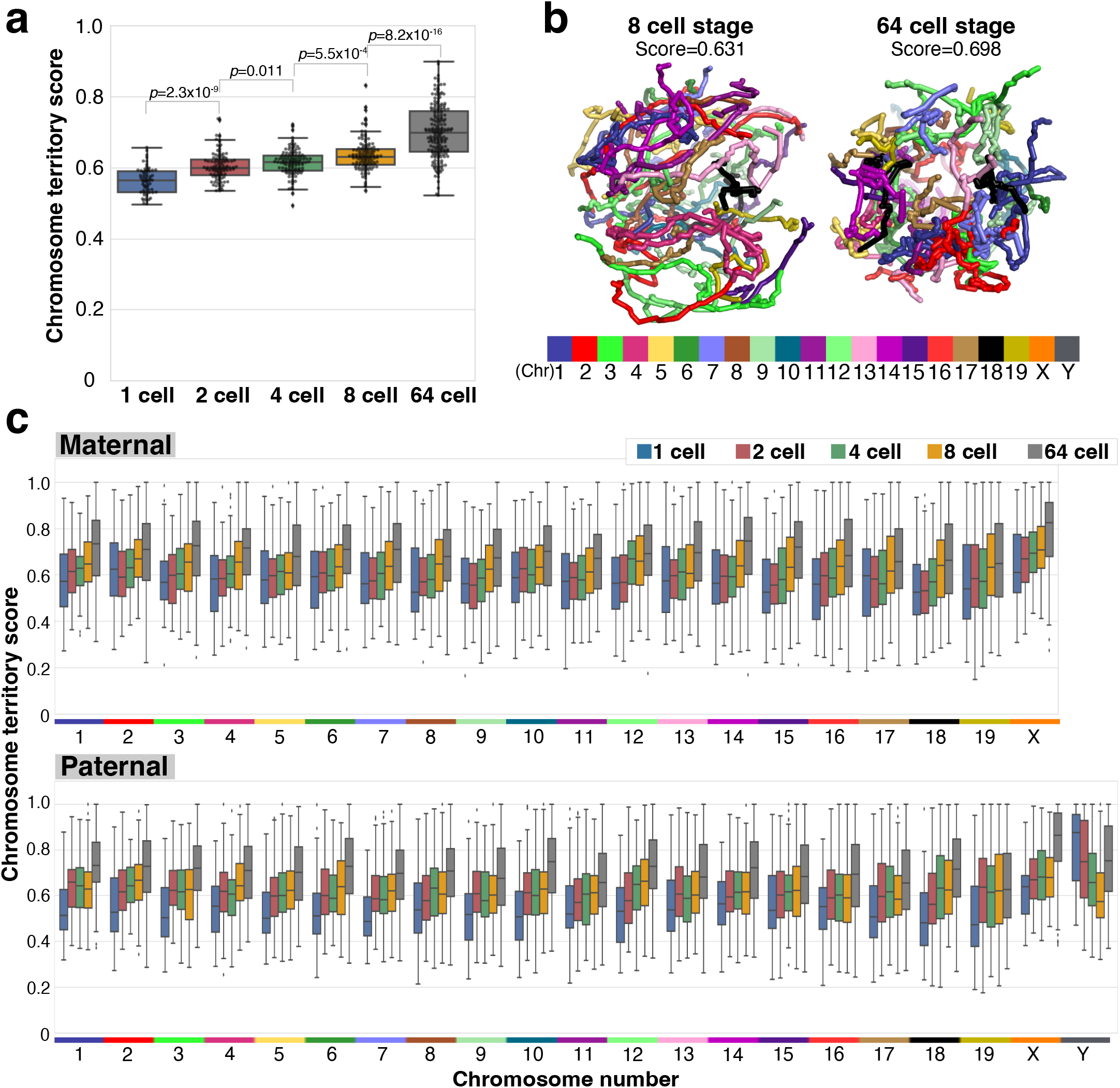
Establishment of discrete chromosome territories during early-stage development. **a**. Box plots showing the chromosome territory score at different developmental stages. Each dot indicates the chromosome territory score of individual cells. The p value was calculated by Mann Whitney U test with Bonferroni correction. **b**. Three-dimensional reconstruction of a representative single nucleus at different developmental stages. Individual chromosomes are labeled in different colors. **c**. Box plots showing the chromosome territory score of each chromosome at different developmental stages.

We also measured the distance between the short- and long-arm termini of each chromosome to assess overall chromosome shape. The average distance between the termini greatly increased from the 2-to 4-cell stages and decreased from the 4-to 64-cell stages (Fig. 4a, b). Greater values of this score appeared to represent an extended (rod-like) shape of chromosomes, found particularly at 4-cell stage (Fig. 4a), and the reduction of this score precedes the onset of discrete (patch-like, nonelongated) CTs from the 8-to 64-cell stages.

**Figure 4:**
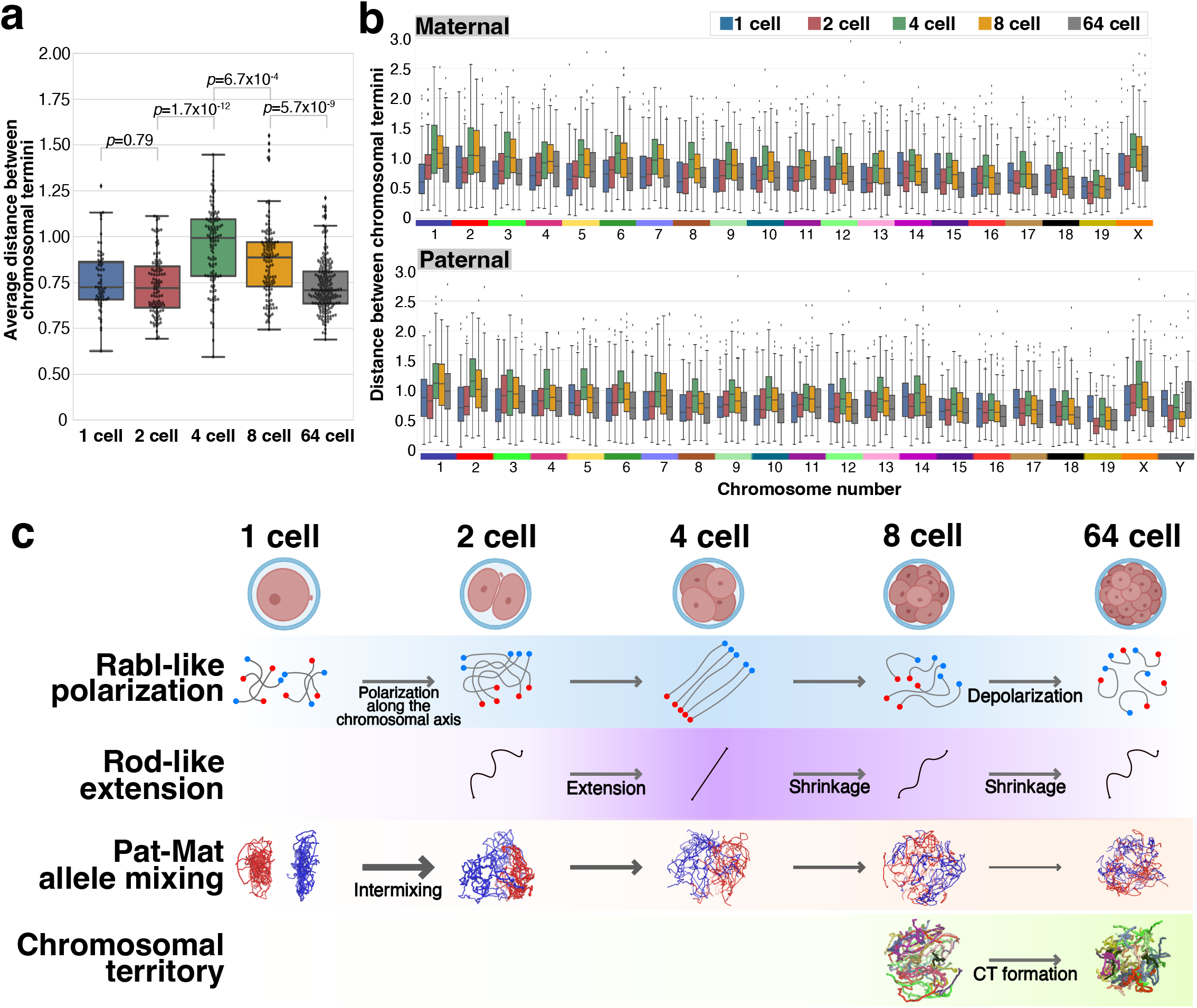
Changes in the chromosomal shape during early-stage development. **a**. Box plots showing the averaged distance between the termini of each chromosome at different developmental stages. **b**. Box plots showing the distance between the termini of each chromosome at different developmental stages. **c**. Schematic summary of chromosomal architecture during early-stage development. Rabl-like polarization of chromosomes starts at the 2-cell stage, which appears to precede rod-like extension and parallel alignment of chromosomes around the 4-cell stage. This rod-like extension is then followed by intermixing of maternal and paternal chromosomes and formation of chromosome territories (CT) especially at the 8-cell and 64-cell stage, respectively. Although causal relationship between these events should be experimentally addressed in future studies, this sequence of events may imply the role of Rabl-like polarization in straightening of chromosomes and effective mixing of chromosomes as suggested in yeast (49, 50), which may facilitate the establishment of chromosome territories by reducing entanglement.

## Discussion

In contrast to bulk Hi-C which can only measure simultaneous contacts between two genomic regions and thus cannot provide information about global genomic organization, the RPR method for scHi-C data can theoretically identify simultaneous contacts among multiple genomic regions and can measure the distance between any genomic regions of interest (even between regions that do not directly contact each other). In this study, we applied the RPR method to construct 3D genomic atlases of early-stage embryos at the single-cell level, with a modification of incorporating the inaccuracy score for data validation, analyzed dynamic changes in global chromosomal architecture at this stage and obtained results that appear to be relevant to the setup process for CT formation including the timing of Rabl-like polarization, rod-like extension and parental allele mixing in relation to CT formation.

Chromosomes are organized into localized CTs that are essential for the higher-order nuclear architecture. It remained unclear, however, when and how CTs are formed at the early stage of embryos. We estimated the extent of CT formation (the CT score) in this study as the proportion of intra-chromosomal interactions, among all chromosomal interactions, between 1-Mb genomic segments as estimated by the RPR method. Our results revealed that discrete CTs are mostly established between 8- and 64-cell stages during early embryonic development. Importantly, this timing appears to be preceded by the dissolvement of Rabl-like polarized configurations and rod-like chromosomal extension, which may play a role in CT formation as discussed below.

The biological meaning of Rabl-like polarized configuration has remained enigmatic. It has been proposed to be just a result from the anaphase orientation of chromosomes without particular cellular roles. Of note, although Rabl-like polarized configuration (which involves separate clustering of telomere and centromere termini) starts to form at 2 cell stage (Fig. 2), the shape of chromosomes starts to elongate only at 4 cell stage (Fig. 4). And both Rabl-like polarization and rod-like extension of chromosomes disappear around the time of CT formation at 64 cell stage. This sequence of events implies a possible role of Rabl-like polarization in rod-like extension and parallel alignment of chromosomes, which may in turn avoid entanglement and thus promote effective mixing of chromosomes, as proposed in yeast as “horsetail movement” (49, 50). The resultant mixing of chromosomes, especially those from different parental origins, may be necessary for establishing proper CTs and therefore should be completed before discrete CTs start to give rise. In addition to mixing of chromosomes, dissolvement of entanglement per se may also facilitate efficient CT formation. This makes sense because establishment of chromosome territories (with little spatial overlap between individual chromosomes) would not be simple if entanglement of chromosomes (inherited from parental gametes) persists.

Interestingly, we found that a fraction of 64 cell stage cells still showed Rabl-like polarization of chromosomes, highlighting cell-to-cell variation of chromosome configuration even among cells at the same developmental stage (Fig. 2a). This variation may reflect the time from mitosis or specific cell types.

Radial positioning of CTs is not random but depends on chromosomal size, GC content, gene density, and transcriptional activity (see refs in Bolzer et al. Plos Biol 2004, Fritz et al. 2019). These factors affect CT positioning in a nuclear shape–dependent and cell type–dependent manner. For example, CT position correlates with gene density in cell types with flat-ellipsoidal nuclei (such as fibroblasts) but not in those with spherical nuclei (such as lymphocytes) (51). Consistent with these observations, we found no significant correlation between gene density and radial CT positioning in early-stage embryonic cells, which have spherical nuclei due to weak cell–cell interactions (Extended Figure 3).

In summary, by analyzing global chromosomal architecture, we observed a sequence of events (Rabl-like polarization and rod-like extension) that may be an important preparatory step for effective chromosomal mixing and subsequent CT formation. In future studies, it would be of great interest to overlay other chromatin-related information onto the 3D genome architecture revealed by the RPR analysis at a single cell level. The RPR analysis in this study was performed at a 1 Mb resolution. Analyses at a higher resolution would be necessary for identification of contacts between gene promoters and enhancers and their relations to epigenetic information such as DNA and histone modifications as well as to TADs and compartments.

## Material and methods

### scHi-C data processing

Processed single cell Hi-C (scHi-C) data of early stage mouse embryos were obtained from Collombet et al. (5) (GSE129029). Details about sequence data processing (i.e., quality control, Hi-C data mapping and allele assignment of sequencing reads) were provided with that data; and detailed information about each result, in supplemental files.

### Reconstruction of 3D chromosome structure from a single diploid cell Hi-C dataset

For each allele of each chromosome, the reference genomic sequence (mm10) was divided into 1-Mb segments. For each segment, a representative point would be prepared in three-dimensional space. Representative points for maternal alleles (from Chr 1 to Chr X) and for paternal alleles (from Chr 1 to Chr X for female cells, from Chr 1 to Chr Y for male cells) were arrayed to form a three-dimensional series so that we can regard this contact map as a recurrence plot.

Assume that a set of phased contacts is given. First, we map this set of phased contacts to the corresponding contact map *C* with the resolution of 1Mb, whose element *C* (*i, j*) is 1 if segments *i* and *j* have at least a phased contact in the set. Otherwise, we set *C* (*i, j*) = 0. In addition, we define *C* (*i, i*) = 1 for every segment *i* for the notational convenience. Therefore, the matrix *C* becomes symmetric, i.e., *C* (*i, j*) = *C* (*j, i*).

Second, we prepare a recurrence plot denoted by a matrix *R* based on the contact map *C*. The matrix *R* is in the same size as *C*. We define *R* (*i, j*) = 1 if there exists a pair of (*k, l*) which corresponds to the same pair of the same alleles for the same chromosomes as (*i, j*), as well as satisfy |*k* − *i*| ≤ 10, |*j* − *l*| ≤ 10, and *C* (*k, l*) = 1. Therefore, the matrix *R* is also symmetric, i.e., *R* (*i, j*) = *R* (*j, i*). This kind of definition for *R* means that the consecutive segments for the corresponding chromosomes are also spatial neighbors, facilitating below the reconstruction of 3D chromosome structure (1, 52).

Third, we construct a network based on *R*. In this network, each segment corresponds to a node in the network. We assign an edge between *i* and *j* if *R* (*i, j*) = 1. For each edge between *i* and *j*, we assign the following local distance *d*:

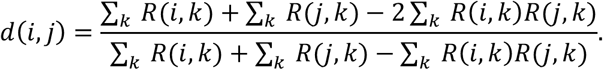

This local distance is the “one-complement” of the Jaccard coefficient (40), which is literally the ratio of unshared neighbors between *i* and *j* against neighbors for either of *i* or *j*. From this third step, we follow Hirata et al. (1, 42).

Fourth, we find the shortest distance for every pair of nodes in this network to obtain a set of global distances. For this step, we here use Johnson’s method (53).

Fifth, we apply the multidimensional scaling to the set of global distances. The multidimensional scaling is a method for finding a set of point arrangements by preserving as a set of distances among given points. We can regard the top three components of the resulting series as our reconstruction of 3D chromosome structure. Let {*x*(*i*) ∈ ℛ^3^} be our reconstruction of 3D chromosome structure.

### Inaccuracy score

We define an inaccuracy score for each reconstruction point *x*(*c*) as follows: First, find the set *N*_*c*_ of the neighboring points in the recurrence plot, i.e. *N*_*c*_ = {*i: R*(*i, c*) = 1}. Then, the inaccuracy score for *x*(*c*) can be defined as

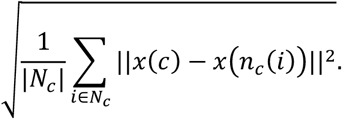

Namely, in this inaccuracy score, we evaluate how spread the neighboring points in the recurrence plot are in our reconstruction.

### Visualization of RPR

We visualized the 3D chromosome structure regarding the top three components obtained from RPR analysis as 3D coordinates. Each segment was visualized in 1 Mb resolution and the overall structure was displayed as beads-on-string structure by Pymol (version 2.4.0) software.

### Exclusion of cell using the inaccuracy score

Standard deviation (s.d.) of the inaccuracy score was calculated to exclude cells with inaccurate structure as depicted in Fig. 1e. We set the threshold of the inaccuracy score s.d. as 0.15 and the cells over this value were removed from further analyses.

### Calculation of maternal-paternal intermixing score

Euclidean distances between each pair of all 1-Mb segments were calculated. The 96th percentile of shortest distances was a threshold for deciding whether any two bins were closely located. A “maternal-paternal intermixing score” for each 1-Mb segment was calculated as the proportion of neighboring segments with different alleles (out of all neighboring segments). Neighboring segments in the same chromosome were excluded from this calculation to remove the bias of genomically close segments. The average score for each cell was determined by averaging the overall maternal-paternal intermixing score of all 1-Mb segments.

### Calculation of mean chromosomal position of proximal pairs

In this analysis, we followed the method of Payne et al. (45) with some modifications. “Chromosomal position” for each 1Mb segment was calculated as the relative position in the chromosomal axis, which took continuous value from 0 to 1 (the short arm terminus = 0, the long arm terminus = 1). Mean chromosomal position of the proximal pairs (“Proximal position score”) was calculated excluding pairs on the same chromosome as the same as the calculation of allele mixing score. Finally, all chromosomes were divided into 100 segments and average proximal position score for each segment was compared with their chromosomal position.

### Calculation of chromosome territory score

Chromosome territory score (CT score) was calculated as the proportion of neighboring segments on the same chromosome out of all neighboring segments. The average score for each cell was determined by averaging the overall CT score of all 1 Mb segments.

### Correlation between gene density and radial CT positioning

Protein coding gene density per 1 Mb for each chromosome was calculated based on comprehensive gene annotation GTF file (https://www.gencodegenes.org/mouse/release_M20.html). Average radial CT positioning for each chromosome at each cell stage was determined by averaging radial CT positioning of all 1 Mb segments of each chromosome.

## Data Availability

Processed single cell Hi-C (scHi-C) data were obtained from Collombet et al. (5) (GSE129029). The processed data from this study are available at Zenodo (https://doi.org/10.5281/zenodo.6658789).

## Author contribution

Conception: Y.K., H.S, Y.G, Y.H., Investigation and data analysis: Y.K. and Y.H, Writing: Y.K., H.S., Y.G and Y.H., Funding acquisition: H.S, Y.G. and Y.H., Supervision: H.S., Y.G. and Y.H.

## Acknowledgments

This study was supported by KAKENHI grants from the Ministry of Education, Culture, Sports, Science, and Technology of Japan and the Japan Society for the Promotion of Science (JP16H06481, JP16H06279, JP16H06479 and JP15H05773 to Y.G., 22K15118 to H.S.) as well as by AMED-CREST of the Japan Agency for Medical Research and Development (JP21gm1310004) by the Uehara Memorial Foundation, and by the International Research Center for Neurointelligence (WPI-IRCN), The University of Tokyo Institutes for Advanced Study.

## Figure Legend

**Extended Figure 1.**
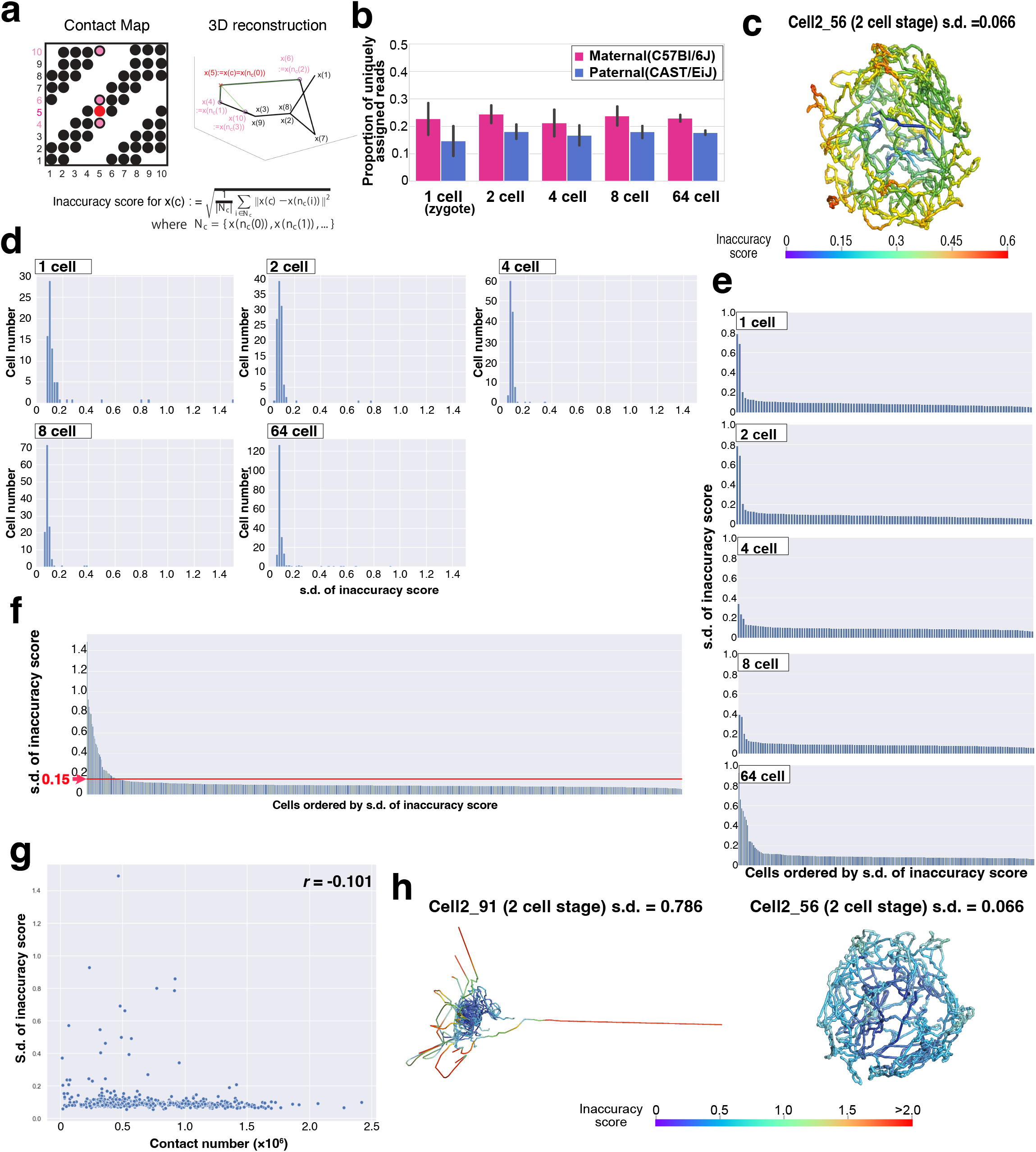
**a**. Schematic illustration of the inaccuracy score. Suppose that we have a contact map on the left side and its 3D reconstruction on the right side. Now we would like to obtain the inaccuracy score for the fifth point shown at the red dot. In the fifth column, there are three other points, which are the fourth point, the sixth point, and the tenth point. These three points are neighbors for the fifth point specified by the contact map. The fifth point is at the red cross on the 3D reconstruction, and the neighbors correspond to the magenta circles. First, we obtain the squared distances of the current fifth point to each of the corresponding neighboring points (fourth, sixth, and tenth points) in the 3D reconstruction. Then, we take their average and calculate its square root, which is the inaccuracy score here. **B**. The proportion of uniquely assigned reads among the total Hi-C contact reads at each cell stage. Standard deviation (s.d.) at each cell stage is shown. Data are represented as mean ± s.d. **c**. The inaccuracy score of Fig. 1d (right) is shown with a different range of the heatmap color. **d**. Bar plots showing s.d. of the inaccuracy score of individual samples at 1 cell, 2 cell, 4 cell, 8 cell and 64 cell stages. The order is sorted by the s.d. of the inaccuracy score. **e**. Histograms showing the number of cells at each range of s.d. of the inaccuracy score at each stage. **f**. The distribution of s.d. of the inaccuracy score of each cell at all stages. The order is sorted by the s.d. of the inaccuracy score. We excluded cells with s.d. greater than 0.15 from the following analyses in this study. **g**. Scatter plots showing the correlation between s.d. of the inaccuracy score and the number of Hi-C contact reads in each cell. **h**. Two examples of three-dimensional reconstruction of a nucleus at 2 cell stage (cell2_91 and cell2_56) by the RPR method. The inaccuracy score is shown by the heatmap color. A cell with a relatively high s.d. of inaccuracy score (left, cell2_91) and a cell with a relatively low s.d. of the inaccuracy score (right, cell2_56) are shown.

**Extended Figure 2.**
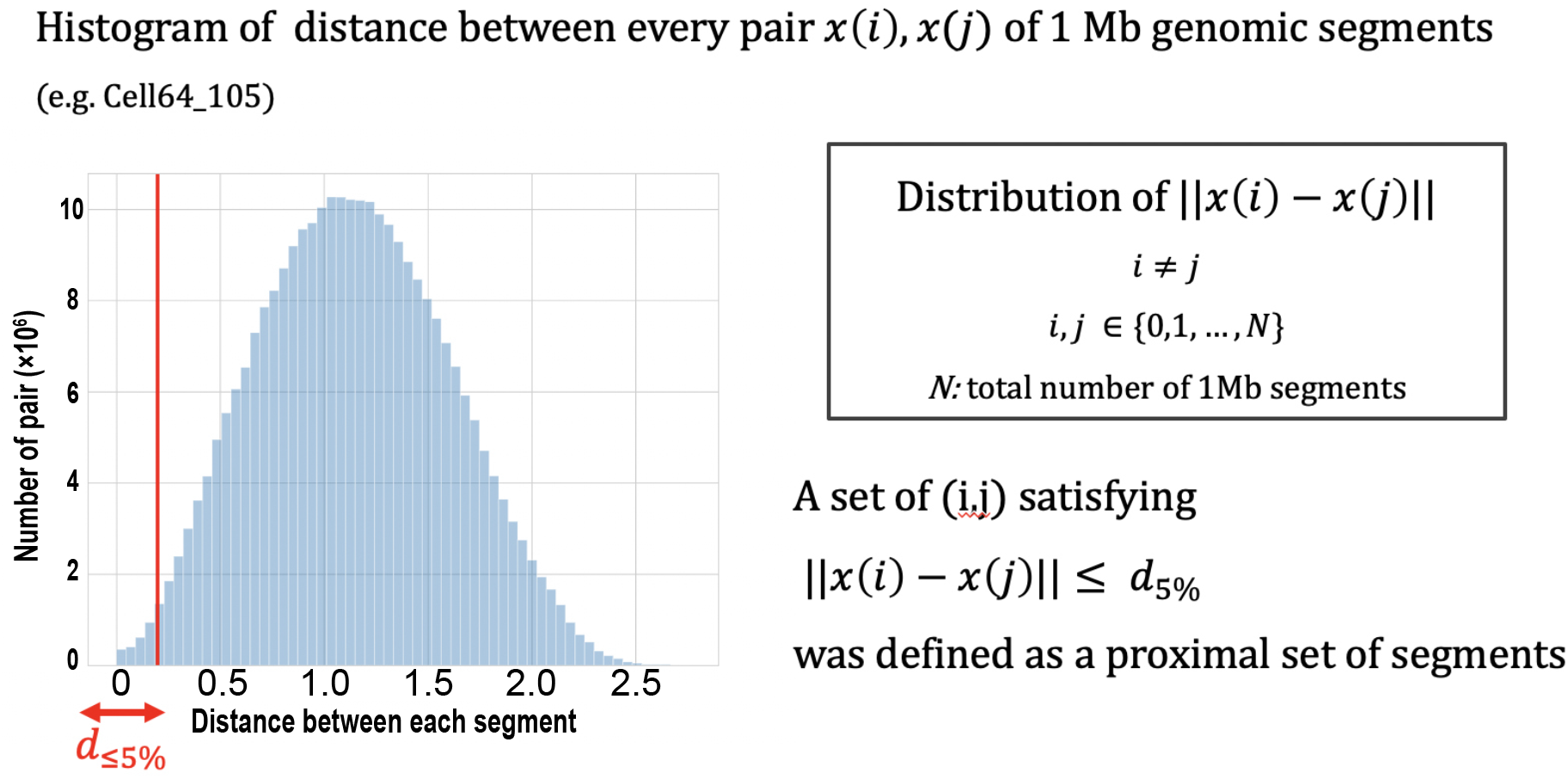
Definition of a proximal set of segments. We defined proximity pairs (the genomic segments located in a proximity) as the top (nearest) 5% of all pairs. These proximity pairs were used for further analyses (Figures 1, 2 and 3).

**Extended Figure 3.**
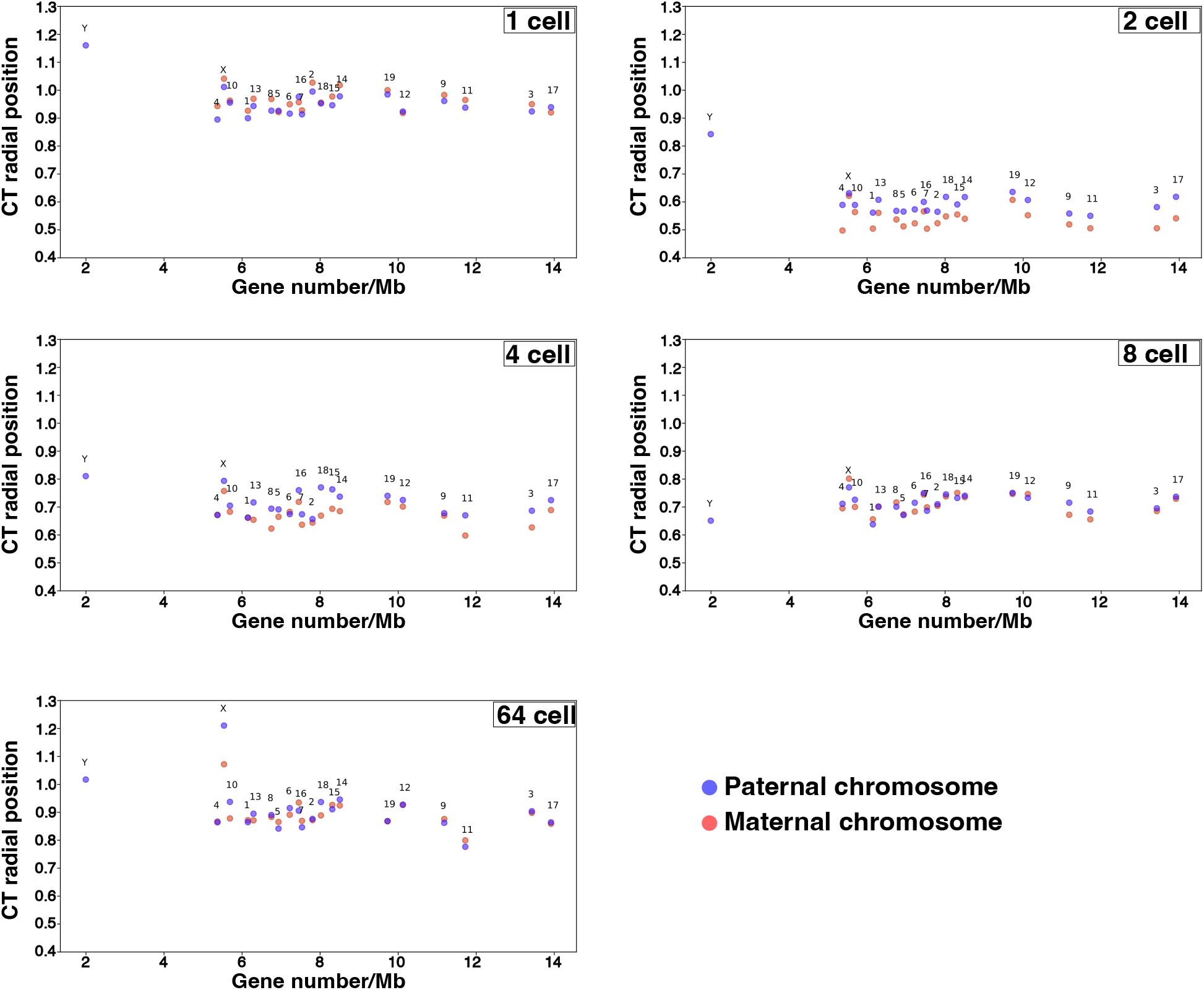
The scatter plot indicates the relationship between gene density and the chromosome territory score (CT score). The CT score was calculated as the proportion of neighboring segments on the same chromosome out of all neighboring segments.

